# Notch signalling maintains Hedgehog responsiveness via a Gli-dependent mechanism during spinal cord patterning in zebrafish

**DOI:** 10.1101/423681

**Authors:** Craig T. Jacobs, Peng Huang

## Abstract

Spinal cord patterning is orchestrated by multiple cell signalling pathways. Neural progenitors are maintained by Notch signalling, whereas ventral neural fates are specified by Hedgehog (Hh) signalling. However, how dynamic interactions between Notch and Hh signalling drive the precise pattern formation is still unknown. We applied the PHRESH (<u>PH</u>otoconvertible <u>RE</u>porter of <u>S</u>ignalling <u>H</u>istory) technique to analyse cell signalling dynamics *in vivo* during zebrafish spinal cord development. This approach reveals that Notch and Hh signalling display similar spatiotemporal kinetics throughout spinal cord patterning. Notch signalling functions upstream to control Hh response of neural progenitor cells. Using gain- and loss-of-function tools, we demonstrate that this regulation occurs not at the level of upstream regulators or primary cilia, but rather at the level of Gli transcription factors. Our results indicate that Notch signalling maintains Hh responsiveness of neural progenitors via a Gli-dependent mechanism in the spinal cord.

## Introduction

Patterning of the spinal cord relies on the action of multiple cell signalling pathways with precise spatial and temporal dynamics (Briscoe and Novitch, 2008). Neural progenitors in the spinal cord are organised into discrete dorsoventral (DV) domains that can be identified by the combinatorial expression of conserved transcription factors (Alaynick et al., 2011; Dessaud et al., 2008; Jessell, 2000). Differentiated post-mitotic neurons migrate from the medial neural progenitor domain to occupy more lateral regions of the spinal cord.

To achieve precise patterning, the developing spinal cord employs anti-parallel signalling gradients of Bone Morphogenic Protein (BMP) and Hedgehog (Hh) to specify dorsal and ventral cell fates, respectively (Le Dréau and Martí, 2012). Cells acquire their fates via sensing both graded inputs. This dual signal interpretation mechanism provides more refined positional information than separate signal interpretation (Zagorski et al., 2017). The action of Sonic Hedgehog (Shh) in the ventral spinal cord is one of the most well studied examples of graded morphogen signalling (Briscoe and Therond, 2013; Cohen et al., 2013). In vertebrates, Hh signalling requires the integrity of primary cilia, microtubule-based organelles present on the surface of most cells (Eggenschwiler and Anderson, 2007). In the absence of the Shh ligand, the transmembrane receptor Patched (Ptc) represses signal transduction by inhibiting the ciliary localisation of the transmembrane protein Smoothened (Smo). When Shh binds to Ptc, Smo inhibition is released, allowing Smo to translocate to the primary cilia (Corbit et al., 2005; Rohatgi et al., 2007). This leads to the activation of the Gli family of transcription factors, resulting in expression of downstream target genes such as *ptc.* Shh thus controls the balance between full-length Gli activators and proteolytically processed Gli repressors (Huangfu and Anderson, 2006; Humke et al., 2010). In mouse, Gli2 is the main activator and its expression does not require active Hh signalling (Bai et al., 2002; Bai and Joyner, 2001). In zebrafish, Gli1 is the main activator (Karlstrom et al., 2003). Although *gli1* is a direct target of Hh signalling, low-level *gli1* expression is maintained in the absence of Hh signalling via an unknown mechanism (Karlstrom et al., 2003). It is thought that Hh-independent *gli* expression allows cells to respond to Hh signals. In the ventral spinal cord, it has been shown that both the level and duration of Hh signalling is critical to the correct formation of the discrete neural progenitor domains along the dorsoventral axis (Dessaud et al., 2010, 2007). However, the temporal dynamics of Hh signalling has been challenging to visualize *in vivo* due to the lack of appropriate tools.

In addition to BMP and Hh signalling, Notch signalling has also been implicated in spinal cord development (Louvi and Artavanis-Tsakonas, 2006; Pierfelice et al., 2011). In contrast to long-range Hh signalling, the Notch signalling pathway requires direct cell-cell interaction, as both receptor and ligand are membrane bound proteins (Kopan and Ilagan, 2009). The Notch receptor, present at the “receiving” cell membrane, is activated by the Delta and Jagged/Serrate family of ligands, present at the membrane of the neighbouring “sending” cell. This leads to multiple cleavage events of Notch, the last of which is mediated by a γ-secretase complex that releases the Notch intracellular domain (NICD). NICD then translocates to the nucleus and forms a ternary transcription activation complex with the mastermind-like (MAML) coactivator and the DNA binding protein RBPJ. This activation complex is essential for the transcription of downstream targets, such as the Hes/Hey family of transcription factors (Artavanis-Tsakonas and Simpson, 1991; Pierfelice et al., 2011). Two major roles of Notch signalling in neural development are to generate binary cell fate decisions through lateral inhibition and to maintain neural progenitor state (Formosa-Jordan et al., 2013; Kageyama et al., 2008). However, how Notch signalling interacts with Hh signalling during spinal cord patterning is not clear.

Using zebrafish lateral floor plate (LFP) development as a model, we previously demonstrated that Notch signalling maintains Hh responsiveness in LFP progenitor cells, while Hh signalling functions to induce cell fate identity (Huang et al., 2012). Thus, differentiation of Kolmer-Agduhr” (KA”) interneurons from LFP progenitors requires the downregulation of both Notch and Hh signalling. Recent reports provide additional support for cross-talk between these pathways during spinal cord patterning in both chick and mouse (Kong et al., 2015; Stasiulewicz et al., 2015). Notch activation causes the Shh-independent accumulation of Smo to the primary cilia, whereas Notch inhibition results in ciliary enrichment of Ptc1. Accordingly, activation of Notch signalling enhances the response of neural progenitor cells to Shh, while inactivation of Notch signalling compromises Hh-dependent ventral fate specification. These results suggest that Notch signalling regulates Hh response by modulating the localisation of key Hh pathway components at primary cilia.

Here, we determine the interaction between Notch and Hh signalling during spinal cord patterning in zebrafish. Using the photoconversion based PHRESH technique, we show that Notch and Hh response display similar spatiotemporal kinetics. Gain- and loss-of function experiments confirm that Notch signalling is required to maintain Hh response in neural progenitors. Surprisingly, Notch signalling doesn’t regulate the Hh pathway at the level of Smo or primary cilia, but rather at the level of Gli transcription factors. Together, our data reveal that Notch signalling functions to control the Hh responsiveness of neural progenitors in a primary cilium-independent mechanism.

## Results

### Generation of a Notch signalling reporter

Spinal cord patterning is a dynamic process with complex interactions of cell signalling pathways in both space and time. To visualise the signalling events in a spatiotemporal manner, we have previously developed the PHRESH (<u>PH</u>otoconvertible <u>RE</u>porter of <u>S</u>ignalling <u>H</u>istory) technique (Huang et al., 2012). This analysis takes advantage of the photoconvertible properties of the Kaede fluorescent protein to visualise the dynamics of cell signalling response at high temporal and spatial resolution. We have utilised the PHRESH technique to visualise Hh signalling dynamics during spinal cord patterning (Huang et al., 2012). To apply the same technique to Notch signalling, we generated a reporter line for *her12*, a target gene of Notch signalling (Bae et al., 2005). This target was chosen because among other Notch target genes co-expressed with *her12*, such as *her2*, *her4*, and *hes5*, *her12* had the highest level of expression throughout the spinal cord (Figure S1). By BAC (bacteria artificial chromosome) recombineering, we generated a *her12:Kaede* reporter by replacing the first coding exon of *her12* with the coding sequence for the photoconvertible fluorescent protein Kaede (Figure 1A). The resulting *her12:Kaede* BAC contains 135 kb upstream and 63 kb downstream regulatory sequences. The *her12:Kaede* reporter line faithfully recapitulated endogenous *her12* expression (Figure 1B). This reporter also responded to different Notch pathway manipulations (Figure 1C-E). The zebrafish *mindbomb* mutant is unable to activate Notch signalling due to an inability to endocytose the Delta ligand (Itoh et al., 2003). As expected, the expression of *her12:Kaede* was completely absent in the spinal cord of *mindbomb* mutants (Figure 1C). Similarly, inhibition of Notch signalling with the small molecule γ-secretase inhibitor LY-411575 (Fauq et al., 2007) completely abolished *her12* expression within 4 hours (Figure 1D and Figure S2). By contrast, ectopic expression of NICD (Notch intracellular domain) using the *hsp:Gal4*; *UAS:NICD* line (Scheer and Campos-Ortega, 1999) resulted in upregulation and expansion of the *her12:Kaede* expression domain (Figure 1E). These results demonstrate that *her12:Kaede* is a sensitive reporter for Notch pathway activity in the spinal cord. The combination of small molecule inhibitors and the *her12:Kaede* reporter allows us to manipulate and monitor Notch signalling dynamics in a tightly controlled temporal manner.

**Figure 1.**
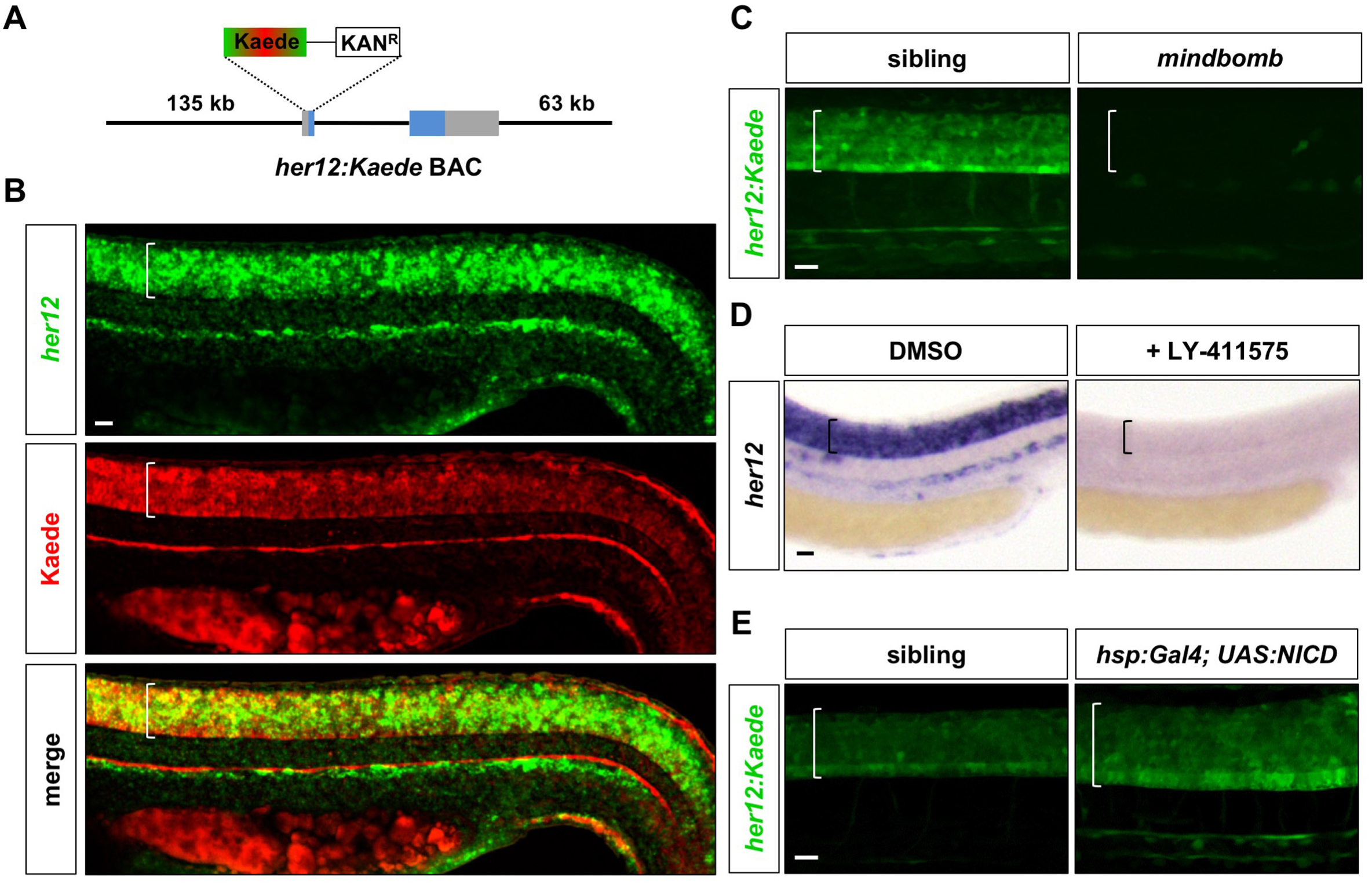
Characterization of the *her12:Kaede* reporter. (A) Schematic drawing of the *her12:Kaede* BAC reporter. A BAC containing the *her12* locus and surrounding regulatory elements was modified to replace the first exon of *her12* with a cassette containing the coding sequence of Kaede and a Kanamycin resistance gene. *her12* coding exons are highlighted in blue. (B) *her12:Kaede* expression, shown by immunohistochemistry using the Kaede antibody (red), recapitulated endogenous *her12* expression, shown by fluorescent *in situ* hybridisation using the *her12* probe (green). (C) *her12:Kaede* expression is completely lost in *mindbomb* mutants at 36 hpf. (D) Inhibition of Notch signalling by LY-411575 from 20 to 30 hpf completely abolished *her12* expression compared to DMSO treated controls. (E) Activation of Notch signalling by *hsp:Gal4*; *UAS:NICD* at 13 hpf resulted in expanded and increased *her12:Kaede* expression at 27 hpf when compared to sibling controls. Brackets in B-E denote the extent of the spinal cord. Scale bars: 20 μm.

### Notch and Hh signalling display similar dynamics during spinal cord patterning

Using the *her12:Kaede* reporter of Notch response (Figure 1) in parallel with the previously described *ptc2:Kaede* reporter of Hh response (Huang *et al*, 2012), we can observe the timing and duration of both pathway activities *in vivo* (Figure 2A). All responding cells are initially labelled by green-fluorescent Kaede (Kaede^green^), which can be photoconverted to red-fluorescent Kaede (Kaede^red^) at any specific time (t_0_). If the cell has finished its signalling response prior to t_0_, only perduring Kaede^red^ will be detected. Conversely, if the cell begins its response after t_0_, only newly synthesised, unconverted Kaede^green^ will be present. Finally, if the cell continuously responds to the signalling both before and after t_0_, a combination of newly-synthesised Kaede^green^ and perduring Kaede^red^ can be observed and the cell will appear yellow. Thus, Kaede^red^ represents “past response” before t_0_, Kaede^green^ indicates “new response” after t_0_, whereas Kaede^red+green^ corresponds to “continued response” through t_0_ (Figure 2A, Video S1). For example, if the embryo is photoconverted at 36 hpf (hours post-fertilization) and imaged 6 hours post-conversion at 42 hpf (36 hpf + 6h), *Kaede^red^* cells have terminated their signalling response before 36 hpf, *Kaede^green^* cells initiate the signalling response between 36 and 42 hpf, while *Kaede^red+green^* cells have sustained signalling response from before 36 hpf and up to a point before 42 hpf. In our experiments, we photoconverted both *ptc2:Kaede* and *her12:Kaede* embryos at 6 hour intervals throughout spinal cord development, and imaged their Kaede fluorescent profiles 6 hours post-conversion of each time point. The time interval of 6 hours was chosen as it allowed time for higher levels of Kaede^green^ to be synthesised while still providing high temporal resolution. We used 3D reconstruction of lateral z-stacks to generate transverse views in order to analyse both the dorsoventral and mediolateral signalling profiles at each time point. Importantly, these signalling profiles were largely similar throughout the anterior-posterior axis of the photoconverted region (Video S2 and S3). Through changing the timing of photoconversion, we were able to create a comprehensive spatiotemporal map of cell signalling dynamics in live embryos (Figure 2B-D).

**Figure 2.**
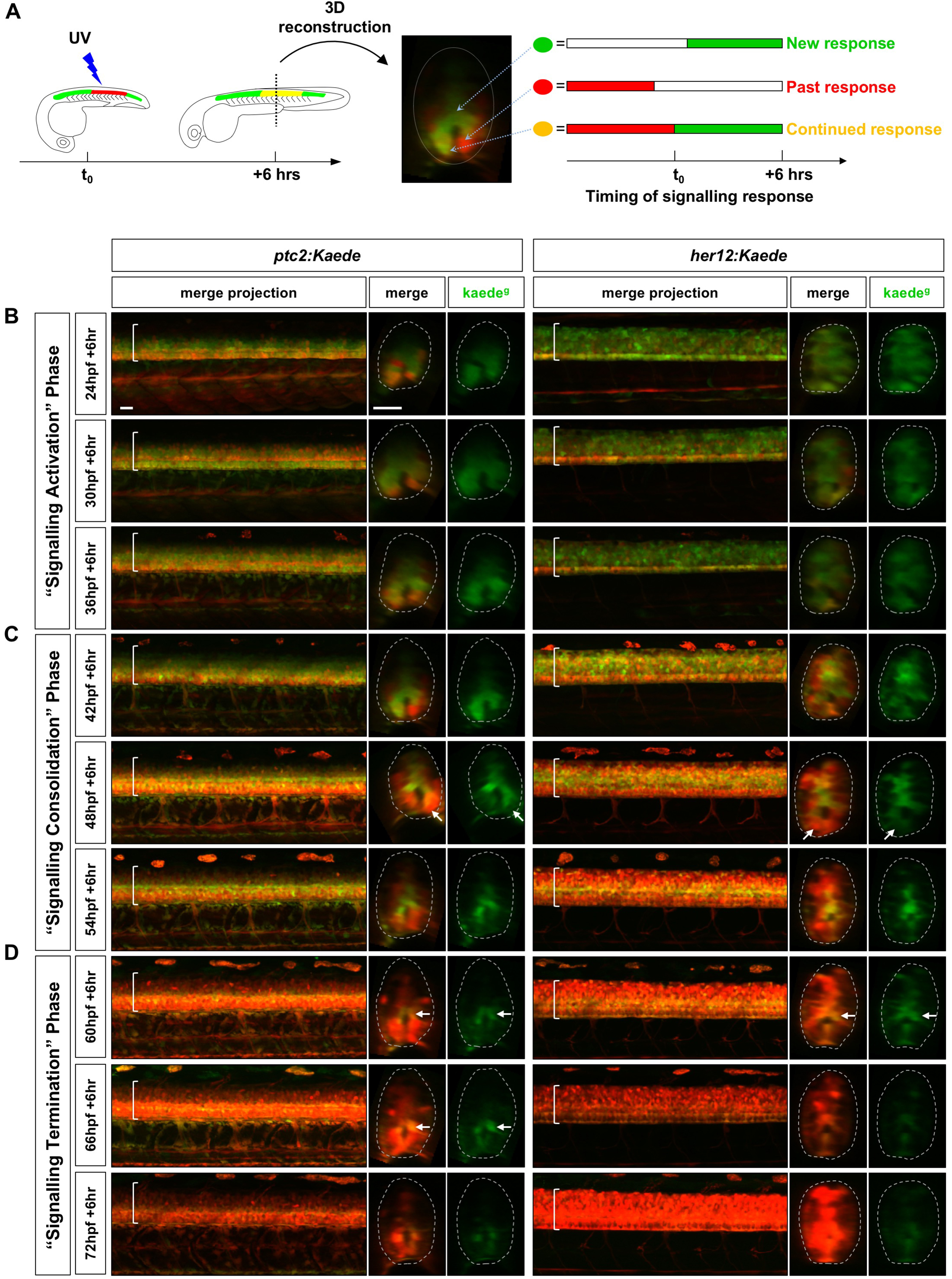
PHRESH analysis of the temporal dynamics of Notch and Hh signalling. (A) Schematic representation of the experimental design. A section of the spinal cord above the yolk extension was photoconverted by the UV light at t_0_ and the fluorescent profile was analysed after 6 hours. 3D reconstruction of the spinal cord allowed the identification of cells that have either new signalling response after t_0_ (green), continued response from before and after t_0_ (yellow), or have ended signalling response before t_0_ (red). (B-D) Time course of Hh and Notch signalling dynamics by PHRESH analysis. *ptc2:Kaede* and *her12:Kaede* embryos were photoconverted at specific time points (t_0_, indicated by hpf) and imaged at 6 hours post photoconversion. Lateral views of confocal projections and transverse views of single slices are shown. Kaede^g^ panels show *de novo* synthesised Kaede^green^ after t_0_, while the merge panels show both previous Kaede^red^ expression and new Kaede^green^ expression. Three distinct phases of signalling response were observed: “signalling activation” phase between 24hpf and 42hpf (B); “signalling consolidation” phase between 42hpf and 60hpf (C); and “signalling termination” phase between 60hpf and 78hpf (D). Arrows in C indicate ventral cells that have terminated response. Arrows in D highlight medial cells right above the spinal canal that remain responsive. Brackets in lateral views and dotted lines in transverse views denote the extent of the spinal cord. Scale bars: 20 μm.

Based on spatiotemporal maps of Notch and Hh response, we divided the signalling dynamics of spinal cord development into three general phases: “signalling activation” phase from 24 to 42 hpf, “signalling consolidation” phase from 42 to 66 hpf, and “signalling termination” phase from 66 to 78 hpf. In the first “signalling activation” phase, active Notch response occurred along the entire dorsoventral axis of the spinal cord (Figure 2B, right), while active Hh response constituted roughly the ventral 75% of the spinal cord in a graded manner (Figure 2B, left). This pattern is consistent with the model that Notch signalling maintains neural progenitor domains, whereas Hh signalling patterns the ventral spinal cord. Interestingly, we found that the signalling response was not entirely homogeneous. In *her12:Kaede* embryos, the majority of cells showed continued Notch response throughout the “signalling activation” phase, but there were some isolated cells in which Kaede expression was completely absent. In *ptc2:Kaede* embryos, some cells had terminated their Hh response (marked by Kaede^red^), while the majority of cells with the same dorsoventral positioning had continued Hh response. The differential Hh response at the same dorsoventral axis is reminiscent of the differentiation of the lateral floor plate domain (Huang et al., 2012).

In the “signalling consolidation” phase (Figure 2C), we observed a dramatic remodelling of the response profiles of both pathways. First, there was an extensive increase in *Kaede^red^* domains for both reporters, indicating the termination of signalling response in these cells. This loss of response was localised to the ventral and lateral regions for Hh signalling (Figure 2C, left) and the ventral, lateral and dorsal regions for Notch signalling (Figure 2C, right). Second, the number of *Kaede^red^* cells increased as the “signalling consolidation” phase progressed. Finally, the active signalling domain consolidated into a tight medial region, which sharpened further to encompass 1 – 2 cell tiers directly dorsal to the spinal canal (Figure 2C, 54hpf + 6h).

Finally, during the “signalling termination” phase (Figure 2D), both active Notch and active Hh responses (Kaede^green^) slowly reduced to the basal level, and most of the spinal cord was marked by Kaede^red^. The active Notch response was notably weaker and restricted to a small medial domain above the spinal canal by 66 hpf before returning to a basal level by 72 hpf. Similarly, active Hh response marked a small medial domain at 66 hpf and 72 hpf, and reduced to a basal level by 78 hpf. After the end of the “signalling termination” phase, both pathways remained at the basal level as spinal cord development progressed (Figure S3).

Comparison of spatiotemporal signalling profiles reveals that Hh and Notch signalling share similar responsive domains. To examine this directly in the same embryo, we performed double fluorescent in situ hybridisation to visualise *her12* and *ptc2* expression together during all three signalling phases of spinal cord development (Figure S4A). During the signalling activation phase (Figure S4A, 24 hpf), *ptc2* expression constituted the ventral portion of the *her12* expression domain, while during the signalling consolidation and termination phases (Figure S4A, 48 hpf and 72 hpf, respectively), *ptc2* and *her12* expression was present within the same restricted medial domain. These results confirm that Notch and Hh response is active within the same cells of the spinal cord. Indeed, double labelling with neural progenitor cell marker *sox2* showed that the medial domain with continued Notch and Hh response corresponded to the *sox2*^+^ neural progenitor domain (Figure S4B-C).

Together, our analysis reveals that Hh signalling response follows similar spatiotemporal kinetics as Notch signalling response during spinal cord patterning, suggesting that Notch signalling plays a role in maintaining Hh response in neural progenitor cells.

### Notch signalling maintains Hh response

To explore the mechanism of interaction between the Notch and Hh signalling pathways, we first performed loss-of-function experiments combining small molecule inhibitors with PHRESH analysis (Figure 3A). We used the Smoothened antagonist cyclopamine (Chen et al., 2002) and the γ-secretase inhibitor LY-411575 to block Hh and Notch signalling, respectively, in a temporally controlled manner. Cyclopamine significantly reduced *ptc2* expression within 4 hours, whereas LY-411575 dramatically downregulated *her12* expression within 2 hours, and completely abolished it after 4 hours of incubation (Figure S2). To ensure complete inhibition of signalling response by the point of photoconversion, *ptc2:Kaede* and *her12:Kaede* embryos were incubated with the inhibitors starting from 20 hpf, photoconverted at 24 hpf and then imaged at 30 hpf, comprising 10 hours of total drug inhibition. As expected, cyclopamine treated *ptc2:Kaede* embryos displayed a marked reduction in the amount of *de novo* synthesised Kaede^green^ compared to controls at 30 hpf (Figure 3A). Similarly, LY-411575 treated *her12:Kaede* embryos showed an almost complete loss of Kaede^green^. In the reciprocal experiments, when *her12:Kaede* embryos were treated with cyclopamine, there was little effect on the levels of Kaede^green^. However, LY-411575 treated *ptc2:Kaede* embryos showed a dramatic reduction in the levels of Kaede^green^, reminiscent of cyclopamine treated *ptc2:Kaede* embryos (Figure 3A). These results suggest that Notch signalling is required for maintaining Hh response, but not vice versa. Interestingly, despite the inhibition of Notch signalling, cells outside of the spinal cord in *ptc2:Kaede* embryos maintained their normal Hh response, indicated by Kaede^green^ expression (Arrowheads in Figure 3A). This result suggests that regulation of Hh response by Notch signalling is tissue specific.

**Figure 3.**
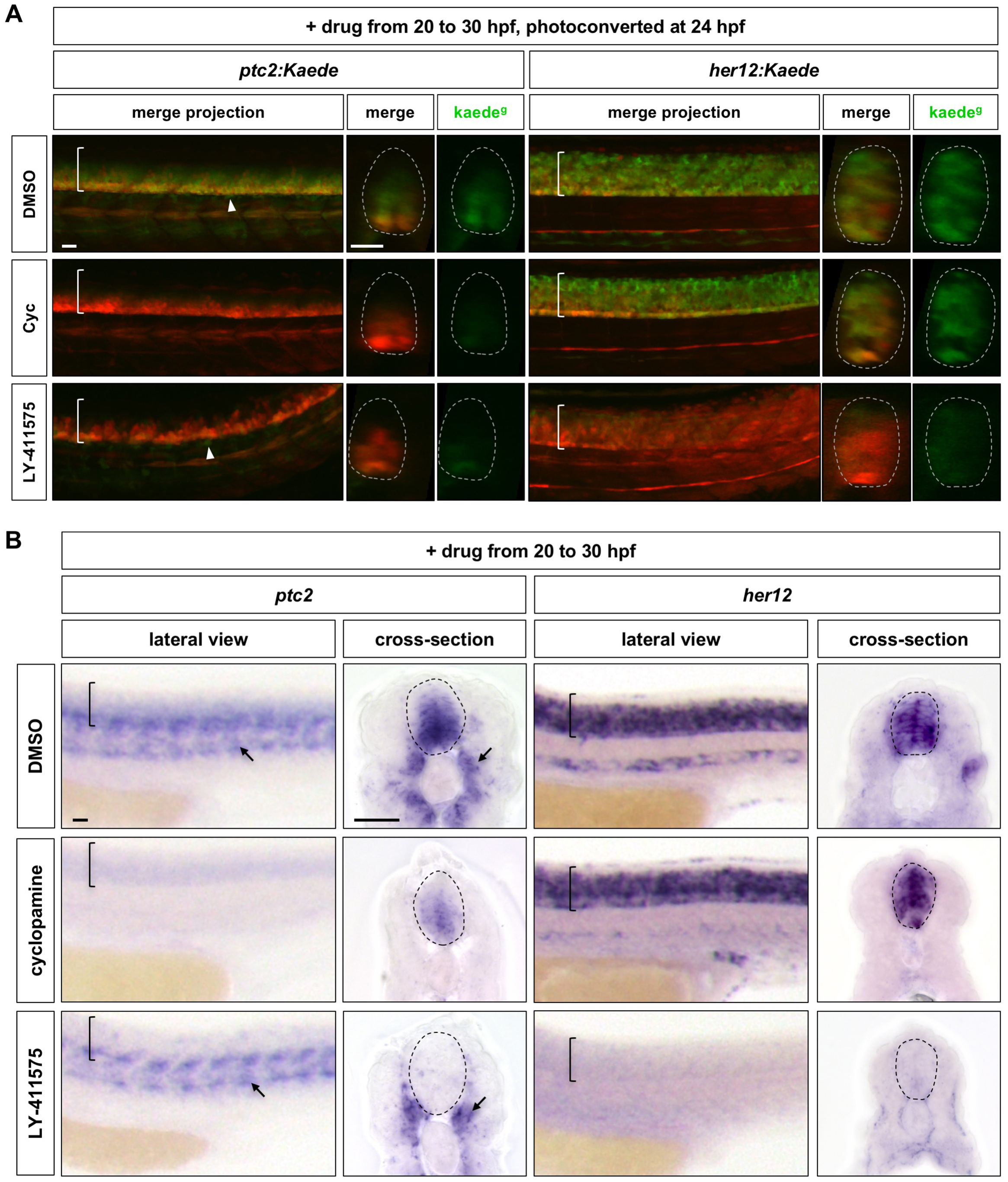
Inhibition of Notch signalling abolishes Hh response in the spinal cord. (A) *ptc2:Kaede* and *her12:Kaede* embryos were incubated with DMSO, cyclopamine (Cyc) or LY-411575 from 20 to 30 hpf, photoconverted at 24 hpf and imaged at 30 hpf. Lateral views of confocal projections and transverse views of single slices are shown. Kaede^g^ panels show *de novo* synthesised Kaede^green^, while the merge panels show both previous Kaede^red^ expression and new Kaede^green^ expression. Arrowheads highlight *Kaede^green^* cells with active Hh response surrounding the notochord. (B) Wild-type embryos were treated with DMSO, cyclopamine, or LY-411575 from 20 to 30 hpf, and stained with *ptc2* or *her12.* Arrows indicate *ptc2* expression in somites. Brackets in lateral views and dotted lines in transverse views in A and B denote the extent of the spinal cord. Scale bars: 20 μm.

To confirm these observations from our PHRESH analysis, we performed RNA in situ hybridisation following small molecule inhibition (Figure 3B). Wild-type embryos were treated with cyclopamine or LY-411575 from 20 to 30 hpf. When Hh signalling was inhibited by cyclopamine, *ptc2* expression in the spinal cord was significantly reduced but not abolished, while *her12* expression in the spinal cord remained unchanged. By contrast, blocking Notch signalling by LY-411575 resulted in complete loss of both *her12* and *ptc2* expression in the spinal cord. As seen in the PHRESH analysis, *ptc2* expression in cells outside of the spinal cord, such as the somites, was largely intact even after Notch inhibition. Similar results were observed in *mindbomb* mutants, where *ptc2* expression was completely abolished in the spinal cord but not in somites (Figure S5), confirming the effect of LY-411575 treatment. Together, these results are consistent with our model that Notch signalling regulates Hh response specifically in the spinal cord. It is also interesting to note that cyclopamine treated embryos showed residual levels of *ptc2* expression in the spinal cord, whereas LY-411575 treatment completely eliminated *ptc2* expression (Figure 3B). It has been shown that zebrafish *smoothened* mutants maintain low-level *gli1* expression in the spinal cord independent of Hh signalling, similar to cyclopamine treated embryos (Karlstrom et al., 2003). The complete loss of *ptc2* expression after Notch inhibition suggests that in contrast to cyclopamine, Notch signalling controls Hh response via a different mechanism, likely downstream of Smo.

In converse experiments, we utilised gain-of-function tools to test the interactions between Notch and Hh signalling (Figure 4). To activate Hh signalling, we used a transgenic line expressing heat shock inducible TagRFP-tagged rSmoM2, a constitutively activate rat Smo mutant (*hsp:rSmoM2*-*TagRFP*). Induction of rSmoM2 at 11 hpf caused a dramatic upregulation of *ptc2* expression throughout the embryo by 24 hpf, but did not affect *her12* expression in the spinal cord when compared to the control (Figure 4A). Interestingly, activation of Hh signalling resulted in a substantial increase in *her12* expression around the dorsal aorta and surrounding vasculature (Figure 4A), suggesting that Hh signalling is upstream of Notch response in the vasculature, the opposite relationship to in spinal cord. To activate ectopic Notch signalling, we induced the expression of the constitutively active Notch intracellular domain (NICD) in *hsp:Gal4*; *UAS:NICD* double transgenic embryos. Induction of NICD at 11 hpf resulted in a spinal cord specific upregulation and expansion of *ptc2* expression at 24 hpf, while the *ptc2* expression pattern external to the spinal cord was unaffected (Figure 4B). Combining our results from the loss- and gain-of-function experiments, we conclude that Notch signalling is required to maintain Hh response, specifically in the spinal cord.

**Figure 4.**
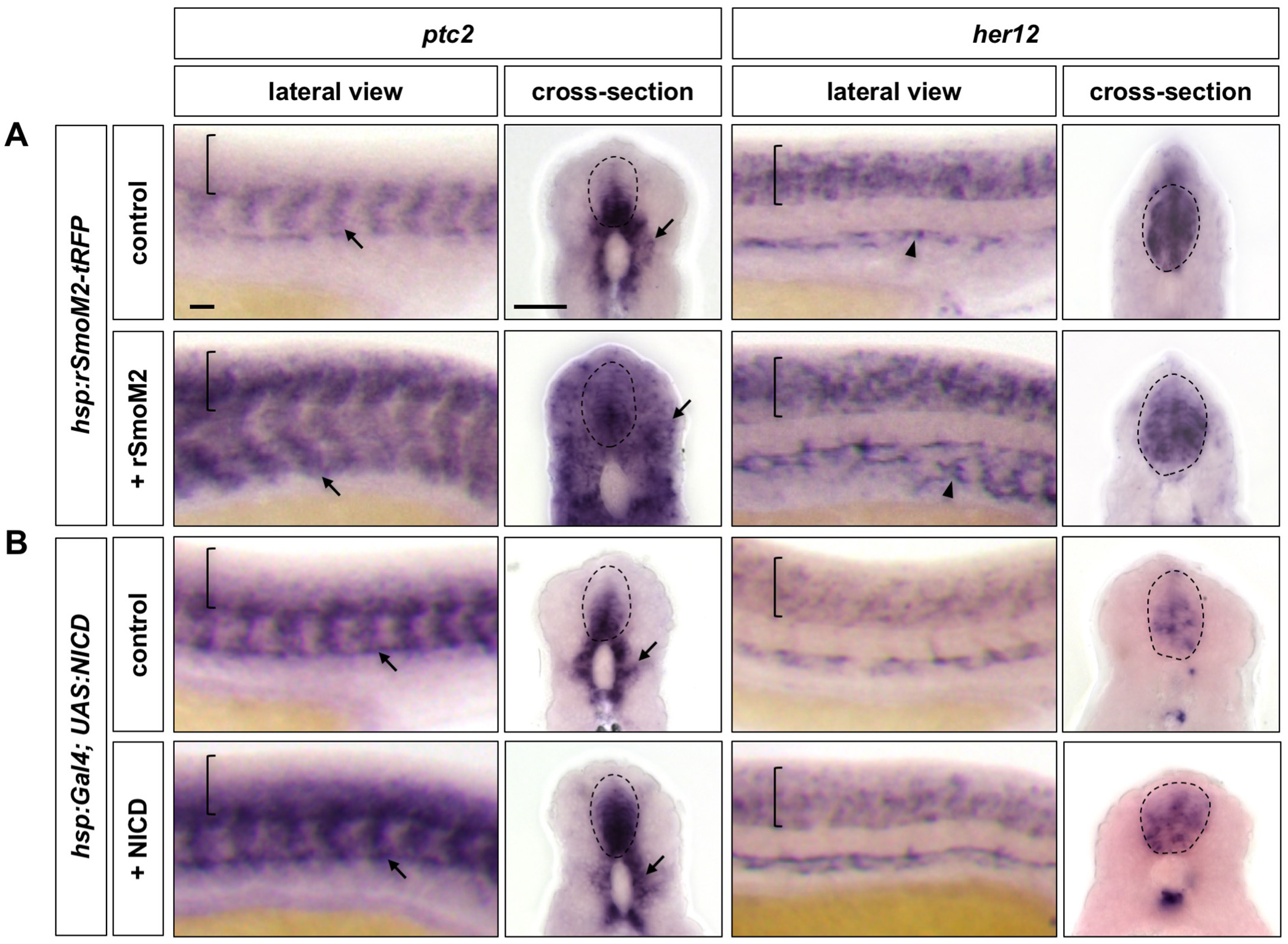
Ectopic Notch activation results in increased and expanded Hh response. *hsp:rSmoM2*-*tRFP* embryos and wild-type controls (A), or *hsp:Gal4; UAS:NICD* embryos and wild-type controls (B) were heat shocked at 11 hpf and stained for the expression of *ptc2* and *her12* at 24 hpf. Brackets in lateral views and dotted lines in transverse views denote the extent of the spinal cord. Arrows indicate *ptc2* expression in somites. Note that expression of *hsp:rSmoM2*-*tRFP* resulted in an expansion of *her12* expression in the vasculature compared to control embryos (arrowheads in A). Scale bars: 20 μm.

### Notch signalling regulates Hh response downstream of Smo

Our results suggest that Notch signalling regulates Hh response during spinal cord patterning. To explore the molecular mechanisms by which Notch controls Hh response, we determined whether activation of Hh signalling at different points of the pathway can bypass the absence of Notch signalling when the small molecule inhibitor LY-411575 is present (Notch^off^ embryos). We first utilised the previously mentioned *hsp:rSmoM2*-*TagRFP* transgenic line to activate Hh signalling at the Smo level (Figure 5A). Wild-type control or *hsp:rSmoM2*-*TagRFP* embryos were heat-shocked at 20 hpf, treated with DMSO or LY-411575 for 10 hours, and assayed for gene expression at 30 hpf (Figure 5B-C). In DMSO treated embryos, induction of rSmoM2 resulted in substantial expansion of *ptc2* expression in both the somites and the spinal cord (Figure 5C). Consistent with this result, activation of Hh signalling by rSmoM2 also led to an expansion of the motor neuron precursor domain, marked by *olig2* expression (Figure 5C). By contrast, when Notch signalling was inhibited by LY-411575, induction of both *ptc2* and *olig2* expression by rSmoM2 was still completely blocked in the spinal cord, similar to LY-411575 treated wild-type controls (Figure 5C). This result suggests that ectopic activation of Hh signalling at the Smo level is not sufficient to restore Hh response in Notch^off^ spinal cords and further implies that Notch signalling likely regulates Hh response downstream of Smo. Interestingly, induction of rSmoM2 did cause an expansion of *ptc2* expression in the surrounding somites despite Notch inhibition (Figure 5C). This observation is consistent with our previous experiments and suggests that this Smo independent mechanism of control is specific to the spinal cord.

**Figure 5.**
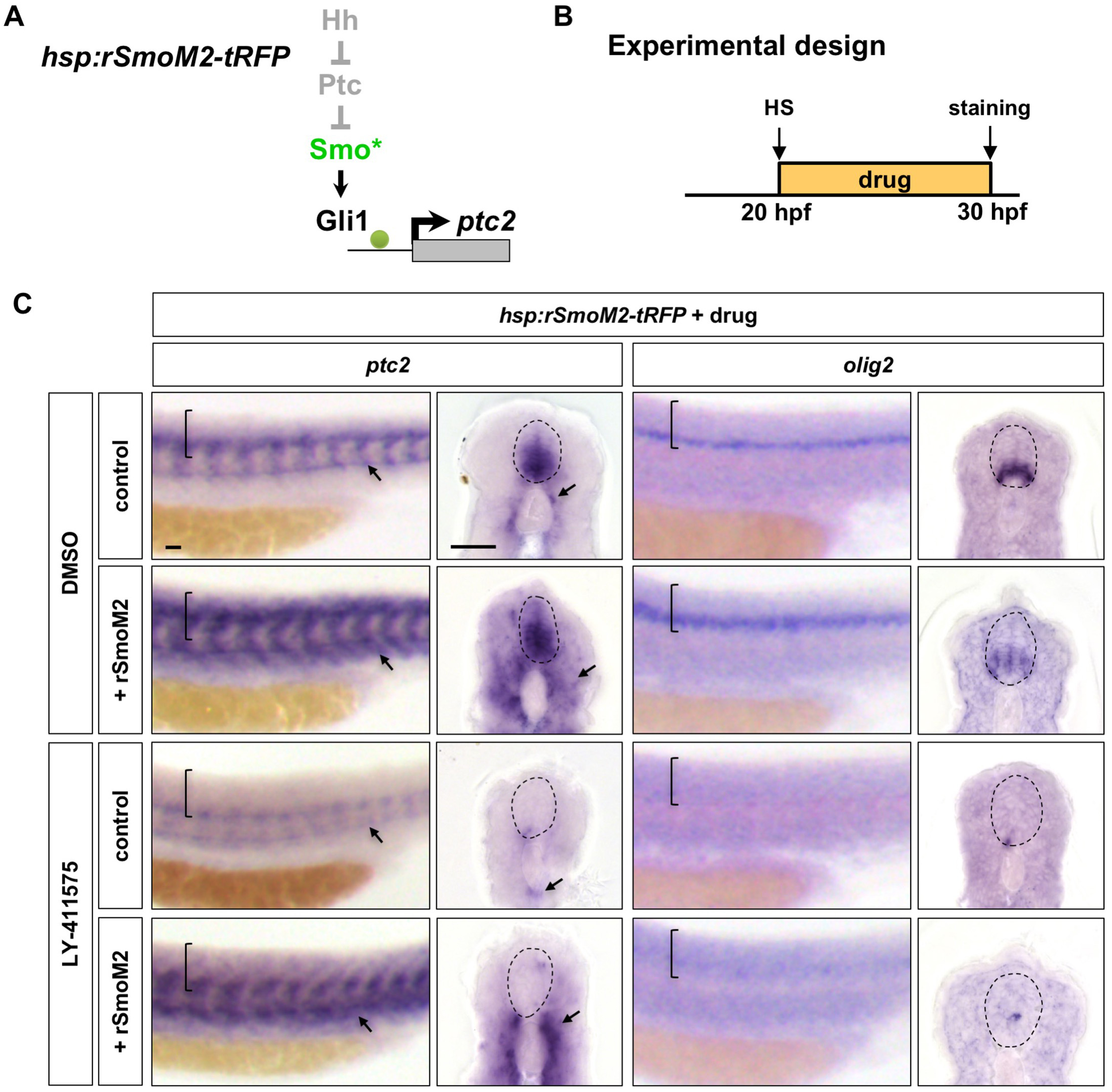
Activation of Hh signalling by rSmoM2 cannot rescue Notch^off^ spinal cords. (A) Schematic representation of the manipulation to the Hh pathway caused by ectopic expression of rSmoM2-tRFP. The point of manipulation is highlighted in green with an asterisk. (B) Experimental design in C. (C) *hsp:rSmoM2*-*tRFP* or wild-type control embryos were heat shocked at 20 hpf, and then incubated in either DMSO or LY-411575 until fixation at 30 hpf. Whole mount in situ hybridisation was performed for *ptc2* and *olig2.* Brackets in lateral views and dotted lines in transverse views denote the extent of the spinal cord. Arrows indicate *ptc2* expression in somites. Scale bars: 20 μm.

### Notch signalling regulates Hh response independent of primary cilia

Vertebrate canonical Hh signalling requires the integrity of primary cilia (Eggenschwiler and Anderson, 2007). To test whether Notch signalling feeds into the Hh pathway via primary cilia, we utilised the *iguana* mutant which lacks primary cilia due to a mutation in the centrosomal gene *dzip1* (Glazer et al., 2010; Huang and Schier, 2009; Kim et al., 2010; Sekimizu et al., 2004; Tay et al., 2010; Wolff et al., 2004). In zebrafish, the complete loss of primary cilia, such as in *iguana* mutants, results in reduction of high-level Hh response concomitant with expansion of low-level Hh pathway activity (Ben et al., 2011; Huang and Schier, 2009). This expanded Hh pathway activation is dependent on low level activation of endogenous Gli1, but does not require upstream regulators of the Hh pathway, such as Shh, Ptc and Smo (Huang and Schier, 2009) (Figure 6A). Thus, the *iguana* mutant also allows us to determine whether low level activation of the endogenous Gli1 transcription factor is able to restore Hh response in Notch^off^ spinal cords. *iguana* mutant embryos or their sibling (heterozygous or wild-type) controls were incubated with DMSO or LY-411575 from 20 hpf and assayed for gene expression at 30 hpf (Figure 6B-C). As shown previously, DMSO treated *iguana* mutants showed a reduction of the highest level of *ptc2* expression in the ventral spinal cord, but displayed an overall expansion of the *ptc2* expression domain in both the spinal cord and somites (Figure 6C). The low level Hh pathway activation in *iguana* mutants was sufficient to induce and expand the *olig2* domain. Remarkably, we found that Hh pathway activation in *iguana* mutants was completely blocked by Notch inhibition (Figure 6C). When *iguana* mutants were treated with LY-411575 at 20 hpf for 10 hours, *ptc2* expression in the spinal cord was completely abolished at 30 hpf, similar to LY-411575 treated sibling controls (Figure 6C). This is in contrast with the somites where *ptc2* expression remained expanded in LY-411575 treated *iguana* mutants similar to DMSO treated *iguana* mutants. Consistent with the loss of *ptc2* expression in the spinal cord, LY-411575 treated *iguana* mutants showed almost no *olig2* expression with only rare scattered *olig2* expressing cells (Figure 6C). Combined with observations from rSmoM2 experiments, these results suggest that Notch signalling likely functions, in a tissue-specific manner, downstream of Smoothened and the primary cilium in its control of Hh response.

**Figure 6.**
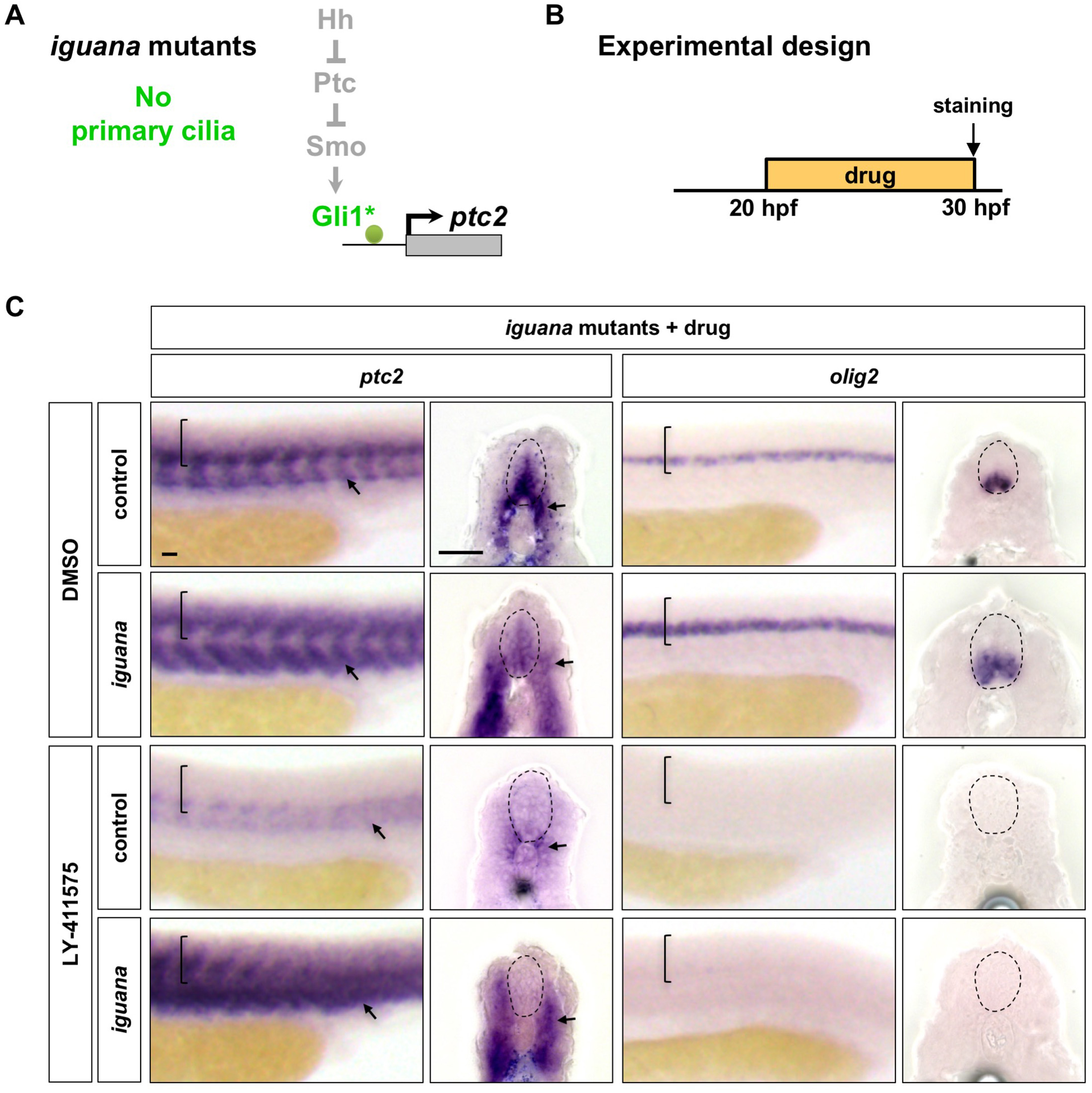
Notch signalling regulates Hh response independent of primary cilia. (A) Schematic representation of the manipulation to the Hh pathway caused by the loss of primary cilia in *iguana* mutants. The point of manipulation is highlighted in green with an asterisk. (B) Experimental design in C. (C) *iguana* mutant and sibling control embryos were incubated in either DMSO or LY-411575 at 20 hpf until fixation at 30 hpf. Whole mount in situ hybridisation was performed for *ptc2* and *olig2.* Brackets in lateral views and dotted lines in transverse views denote the extent of the spinal cord. Arrows indicate *ptc2* expression in somites. Scale bars: 20 μm.

### Ectopic expression of Gli1 partially rescues Hh response in Notch^off^ spinal cords

Since low level activation of endogenous Gli1 in *iguana* mutants is not sufficient to restore Hh response in Notch^off^ spinal cords, we hypothesised that Notch signalling regulates Hh response by maintaining *gli1* expression. To test this possibility, we treated wild-type embryos with DMSO, cyclopamine, or LY-411575 at 20 hpf for 10 hours, then assayed for *gli1* gene expression at 30 hpf (Figure 7A). In DMSO treated controls, *gli1* expression was present throughout the ventral spinal cord and in the somites. In cyclopamine treated embryos, *gli1* expression was dramatically reduced, but a low level remained in the spinal cord, corresponding to Hh-independent *gli1* transcription (Karlstrom et al., 2003). By contrast, LY-411575 treatment completely abolished *gli1* expression in the spinal cord. Strikingly, *gli1* expression in surrounding tissues remained largely unaffected. These results suggest that Notch signalling is required to maintain Hh-independent *gli1* expression in the spinal cord. Interestingly, similar experiments demonstrated that expression of other members of the *gli* genes, *gli2a*, *gli2b* and *gli3*, in the spinal cord, was also largely abolished by LY-411575 treatment (Figure S6), suggesting that Notch signalling controls the expression of all Gli transcription factors.

**Figure 7.**
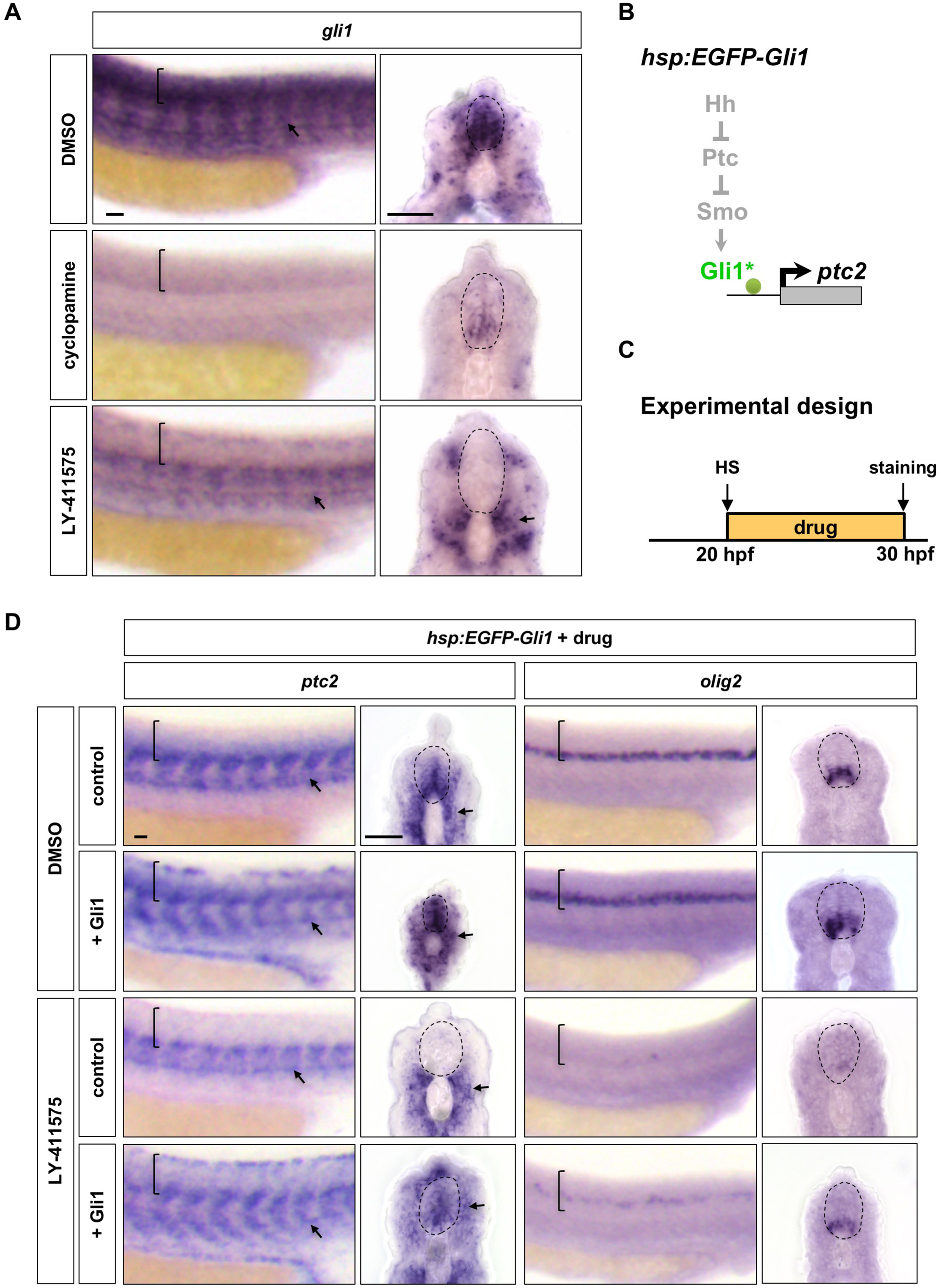
Notch signalling regulates Hh response at the Gli level. (A) Whole-mount in situ hybridisation for *gli1* was performed on wild-type embryos treated with DMSO, cyclopamine, or LY-411575 from 20 to 30 hpf. (B) Schematic representation of the manipulation of the Hh pathway caused by ectopic EGFP-Gli1 expression. The point of manipulation is highlighted in green with an asterisk. (C) Experimental design in D. (D) *hsp:EGFP*-*Gli1* and wild type control embryos were heat shocked at 20 hpf, and then incubated in either DMSO or LY-411575 until fixation at 30 hpf. Whole mount in situ hybridisation was performed for *ptc2* and *olig2.* Brackets in lateral views and dotted lines in transverse views in A and D denote the extent of the spinal cord. Arrows in A and D indicate *ptc2* expression in somites. Scale bars: 20 μm.

We next examined whether overexpression of ectopic Gli1 was sufficient to rescue Hh response in Notch^off^ spinal cords (Figure 7B). We used an EGFP-Gli1 transgene under the control of a heat shock inducible promoter (*hsp:EGFP*-*Gli1*) (Huang and Schier, 2009). Similar to previous experiments, wild type control or *hsp:EGFP*-*Gli1* embryos were heat shocked at 20 hpf, treated with DMSO or LY-411575 for 10 hours, and assayed for gene expression at 30 hpf (Figure 7C-D). Induction of ectopic EGFP-Gli1 resulted in the *ptc2* expression domain expanding further dorsally and throughout the spinal cord and the *olig2* domain was also larger (Figure 7D), a similar phenotype to rSmoM2 induction in DMSO treated embryos. Strikingly, when EGFP-Gli1 induction was followed by LY-411575 treatment, we observed significant *ptc2* expression in the spinal cord, although at a slightly lower level compared to DMSO treated *hsp:EGFP*-*Gli1* embryos (Figure 7D). Critically, the restored Hh pathway activation in LY-411575 treated *hsp:EGFP*-*Gli1* embryos was able to activate *olig2* expression, although substantially weaker than the wild-type level (Figure 7D). Together, these results suggest that, in the spinal cord, Notch signalling regulates Hh response by modulating the Gli1 transcription factor, as ectopic Gli1 can partially rescue the Hh response in Notch^off^ spinal cords. This regulation is partly through transcriptional control of *gli1* expression. However, since the ectopic EGFP-Gli1 was unable to rescue the highest level of Hh response and cannot fully restore *olig2* expression in Notch^off^ spinal cords, it is likely that Notch signalling also plays additional roles in regulating Gli1 protein – such as its post-translational modification or stability.

## Discussion

We provide *in vivo* evidence for cross-talk between two conserved developmental signalling pathways, Notch and Hh signalling, in the zebrafish spinal cord. Through the PHRESH technique, we observe shared spatiotemporal dynamics of pathway activity throughout spinal cord patterning, highlighting a role for Notch and Hh interaction in neural progenitor maintenance and specification. Using both gain- and loss-of function techniques, we establish a primary cilia-independent mechanism by which Notch signalling permits neural progenitors to respond to Hh signalling via *gli* maintenance.

### Studying cell signalling dynamics using PHRESH

We have previously developed the PHRESH technique to study the dynamics of Hh signalling *in vivo* (Huang et al., 2012). In this study, we demonstrate the versatility of the PHRESH method by correlating the dynamics of Hh and Notch signalling *in vivo* using the *ptc2:Kaede* reporter and a new *her12:Kaede* reporter. Traditional transcriptional GFP reporters fail to provide temporal information due to GFP perdurance, whereas destabilised fluorescent protein reporters can only provide current activity at the expense of signalling history. By contrast, the PHRESH technique utilises Kaede photoconversion to delineate the cell signalling history in any given time window by comparing newly synthesised Kaede^green^ (new signalling) with photoconverted Kaede^red^ (past signalling). We envision that PHRESH analysis could be combined with cell transplantation and time-lapse imaging to simultaneously analyse cell lineage and signalling dynamics at single cell resolution. Similar approaches can easily be adapted to study other dynamic events by using photoconvertible fluorescent reporters.

### Spatiotemporal dynamics of Hh and Notch signalling

Using the PHRESH technique, we created a spatiotemporal map of signalling dynamics for the Hh and Notch pathways during spinal cord patterning. Strikingly, Notch and Hh signalling display similar activity profiles. We have characterised these profiles into three general phases: “signalling activation”, “signalling consolidation”, and “signalling termination”. In the early “signalling activation” phase, Notch signalling is active throughout the spinal cord, while active Hh response occurs in the ventral ~75% of the spinal cord. During “signal consolidation”, the responsive domain of both pathways sharpens into a small medial domain dorsal to the spinal canal; in “signalling termination” the response to both pathways returns to a basal level. Our detailed time course reveals three key features of Notch and Hh signalling dynamics. First, early active Hh signalling shows a graded response with the highest level in the ventral domain, as predicted by the classical morphogen model (Briscoe and Small, 2015). By contrast, active Notch response does not appear to be graded along the ventral-dorsal axis. Second, despite showing the highest level of Hh response early, the ventral spinal cord terminates Hh response earlier than the more dorsal domains. Therefore, the ventral domain shows higher level Hh response for a shorter duration, whereas the dorsal domain shows lower level response for a longer duration. Our observation is reminiscent of the floor plate induction in chick and mouse embryos, where the specification of the floor plate requires an early high level of Hh signalling and subsequent termination of Hh response (Ribes et al., 2010). Our result suggests that Hh signalling dynamics is also evolutionarily conserved. Third, lateral regions of the spinal cord lose both Notch and Hh response before the medial domains. As the active signalling response consolidates into the medial domain, so does the expression of *sox2*, a neural progenitor marker, suggesting that neural differentiation is accompanied by the attenuation of Notch and Hh response. Our observation is consistent with the notion that neural progenitors occupy the medial domain of the spinal cord and that as they differentiate they move laterally.

### Notch signalling regulates Hh response

The loss of Hh response is a necessary step for fate specification, as shown in the chick studying floor plate induction (Ribes et al., 2010) and post-mitotic motor neuron precursors (Ericson et al., 1996). We have previously shown that the time at which cells attenuate their Hh response is crucial for fate specification in the zebrafish ventral spinal cord (Huang et al., 2012). How do neural progenitor cells in the spinal cord maintain their Hh responsiveness until the correct time in order to achieve their specific fates? Three independent lines of evidence indicate that Notch signalling is likely part of this temporal attenuation mechanism controlling Hh responsiveness. First, PHRESH analysis reveals that active Hh response correlates with Notch signalling activity. The active Hh signalling domain initially constitutes part of the active Notch response domain before following similar kinetics in signalling consolidation and termination. This result is consistent with the model that Notch signalling is necessary to maintain Hh responsiveness. Second, loss of Notch signalling either by genetic mutants or by small molecule inhibition results in loss of active Hh response in the spinal cord. In contrast, inhibition of Hh signalling does not affect Notch pathway activity. Instead, constitutive activation of Notch signalling leads to enhanced Hh pathway activation. Together, these results suggest that Notch signalling functions upstream of Hh signalling in controlling Hh responsiveness during spinal cord patterning.

How does Notch signalling control Hh response? Previous reports have implicated Notch signalling in the regulation of ciliary trafficking of Smo and Ptc (Kong et al., 2015; Stasiulewicz et al., 2015), thereby modulating cellular responsiveness to Hh signals. By contrast, we show that in the absence of primary cilia in *iguana* mutants, the low level Hh response remaining due to constitutive Gli1 activation can be completely blocked by Notch inhibition. This result suggests that Notch signalling can regulate Hh response in a primary cilium independent manner. In zebrafish, Gli1 functions as the main activator downstream of Hh signalling, although Gli2a, Gli2b and Gli3 also contribute to the activator function (Karlstrom et al., 2003; Ke et al., 2008; Tyurina et al., 2005; Vanderlaan et al., 2005; Wang et al., 2013). Indeed, inhibition of Notch signalling abolishes both Hh-dependent and Hh-independent *gli1* expression in the spinal cord. Similarly, *gli2a*, *gli2b* and *gli3* expression in the spinal cord is largely eliminated in Notch^off^ spinal cords. These results demonstrate that Notch signalling controls the transcription or mRNA stability of all members of the Gli family in the spinal cord. It is possible that *gli* genes are direct targets of Notch signalling, as shown in mouse cortical neural stem cells where N1ICD/RBPJ-κ binding regulates *Gli2* and *Gli3* expression (Li et al., 2012). Importantly, while ectopic expression of Gli1 from the *hsp:EGFP*-*Gli1* transgene can re-establish Hh response as indicated by *ptc2* expression in the Notch^off^ spinal cord, it is unable to fully restore the *olig2* motor neuron precursor domain. This finding argues that Notch signalling likely plays additional roles in regulating Gli1 protein level or activity. A similar mechanism has been suggested in Müller glia of the mouse retina, where Notch signalling controls Gli2 protein levels and therefore Hh response (Ringuette et al., 2016). Interestingly, the study by Ringuette et al. favours a translation or protein stability model because Notch manipulation does not alter the *Gli2* transcript level. Our work demonstrates that Notch signalling functions to permit neural progenitors to respond to Hh signalling via *gli* transcriptional regulation and Gli protein maintenance. It is conceivable that Notch signalling regulates the Hh pathway at the level of both Ptc/Smo ciliary trafficking and the Gli transcription factors. This dual regulation might ensure efficient termination of Hh response during neuronal differentiation. Intriguingly, the regulation of Hh response by Notch signalling appears to be specific to the neural tissue. The Hh response in the somites is largely unaffected by Notch manipulations, whereas activation of Hh signalling results in an expansion of *her12* expression in the blood vessels, suggesting Notch response is likely downstream of Hh signalling in the vasculature.

In summary, we demonstrate that Notch and Hh signalling share similar spatiotemporal kinetics during spinal cord patterning and that this dynamic interaction is likely required to maintain the neural progenitor zone. We also provide evidence for a primary cilium-independent and Gli-dependent mechanism in which Notch signalling permits these neural progenitors to respond to Hh signalling.

## Materials and Methods

### Zebrafish Strains

All zebrafish strains used in this study were maintained and raised under standard conditions and all procedures were approved by the University of Calgary Animal Care Committee. The transgenic strains used were: *ptc2:Kaede* (Huang et al., 2012), *her12:Kaede*, *hsp:Gal4* (Scheer and Campos-Ortega, 1999), *UAS:NICD* (Scheer and Campos-Ortega, 1999), *hsp:rSmoM2*-*TagRFP*, *hsp:EGFP*-*Gli1* (Huang and Schier, 2009). The *hsp:rSmoM2*-*TagRFP* line was generated by standard Tol2-mediated transgenesis. The *mib^ta52b^* (Itoh et al., 2003) and *igu^ts294^* (Sekimizu et al., 2004; Wolff et al., 2004) mutant strains were maintained as heterozygotes and homozygous embryos were generated by intercrossing heterozygous carriers.

### Generation of the *her12:Kaede* BAC transgenic line

To generate the *her12:Kaede* transgenic line, BAC clone zK5I17 from the DanioKey library that contains the *her12* locus and surrounding regulatory elements was selected for bacteria-mediated homologous recombination following standard protocol (Bussmann and Schulte-Merker, 2011). zK5I17 contains 135 kb upstream and 63 kb downstream regulatory sequences of *her12.* First, an iTol2-amp cassette containing two Tol2 arms in opposite directions flanking an ampicillin resistance gene was recombined into the vector backbone of zK5I17. Next, a cassette containing the Kaede open reading frame and the kanamycin resistance gene was recombined into the zK5I17-iTol2-amp to replace the first exon of the *her12* gene. Successful recombinants were confirmed by PCR analysis. The resulting *her12:Kaede* BAC was co-injected with *tol2* transposase mRNA into wild-type embryos and stable transgenic lines were established through screening for Kaede expression.

### In situ hybridisation and immunohistochemistry

All whole-mount in situ hybridisation and immunohistochemistry in this study were performed using standard protocols. We used the following antisense RNA probes: *gli1*, *gli2a*, *gli2b*, *gli3*, *her2*, *her4*, *her12*, *hes5*, *olig2*, *ptc2* and *sox2.* For double fluorescent in situ hybridisation, both dinitrophenyl (DNP) and digoxigenin (DIG) labelled probes were used with homemade FITC and Cy3 tyramide solutions (Vize et al., 2009). For immunohistochemistry, rabbit polyclonal antibody to Kaede (1:1000, MBL) was used. The appropriate Alexa Fluor-conjugated secondary antibodies were used (1:500, Thermo Fisher) for fluorescent detection of antibody staining and Draq5 (1:10,000, Biostatus) was used for nuclei staining.

### Photoconversion for PHRESH analysis

All fluorescent imaging was carried out using an Olympus FV1200 confocal microscope and Fluoview software. Photoconversion was carried out using the 405nm laser with a 20x objective. *ptc2:Kaede* and *her12:Kaede* embryos at the appropriate stages were anaesthetised with 0.4% tricaine and then embedded in 0.8% low melting agarose. To achieve complete conversion over a large area, a rectangular area of 1000 by 300 pixels was converted by scanning the area twice with 50% 405nm laser at 200 μs per pixel. Following confirmation of Kaede^red^ expression, embryos were recovered in E3 water with phenylthiourea for 6 hours post-conversion before imaging. Appropriate imaging parameters were established using the unconverted region as a reference to avoid over or under exposure of the Kaede^green^ signal. Cross-sections were generated using Fiji-ImageJ software (Schindelin et al., 2012) to create a 3D reconstruction of the image, then “resliced” to yield transverse views of the spinal cord.

### Drug treatment

Embryos at the appropriate stage were treated with cyclopamine (Toronto Chemical, 100 μM), LY-411575 (Sigma, 50 μM), or DMSO control in E3 fish water. For PHRESH analysis, embryos were treated from 4 hours prior to the point of conversion until 6 hours post-conversion. To match this, all other drug treatments took place between 20 hpf and 30 hpf.

### Heat shock experiments

To induce expression from the heat shock promoter, embryos at the relevant stage were placed in a 2 ml micro-centrifuge tube in a heat block set to 37°C for 30 minutes. After heat shock, embryos were transferred back into E3 water in a petri dish and recovered at 28.5°C. For drug treatment after heat shock, embryos were transferred directly from the heat shock to E3 water containing the appropriate drug.

### Cryosectioning

To obtain transverse sections after whole-mount in situ hybridisation, embryos were cryoprotected with 30% sucrose at 4°C before being embedded in OCT compound (VWR) and frozen in the −80°C freezer. Sections were cut between 10-16μm using a Leica cryostat. Sections were taken from the region of the trunk dorsal to the yolk extension.

## Acknowledgements

We thank the zebrafish community for providing probes and reagents; Holger Knaut for BAC clones; Sarah Childs and members of the Huang laboratory for discussion; and Paul Mains and James McGhee for critical comments on the manuscript.

## Supplemental Figures

**Figure S1.**
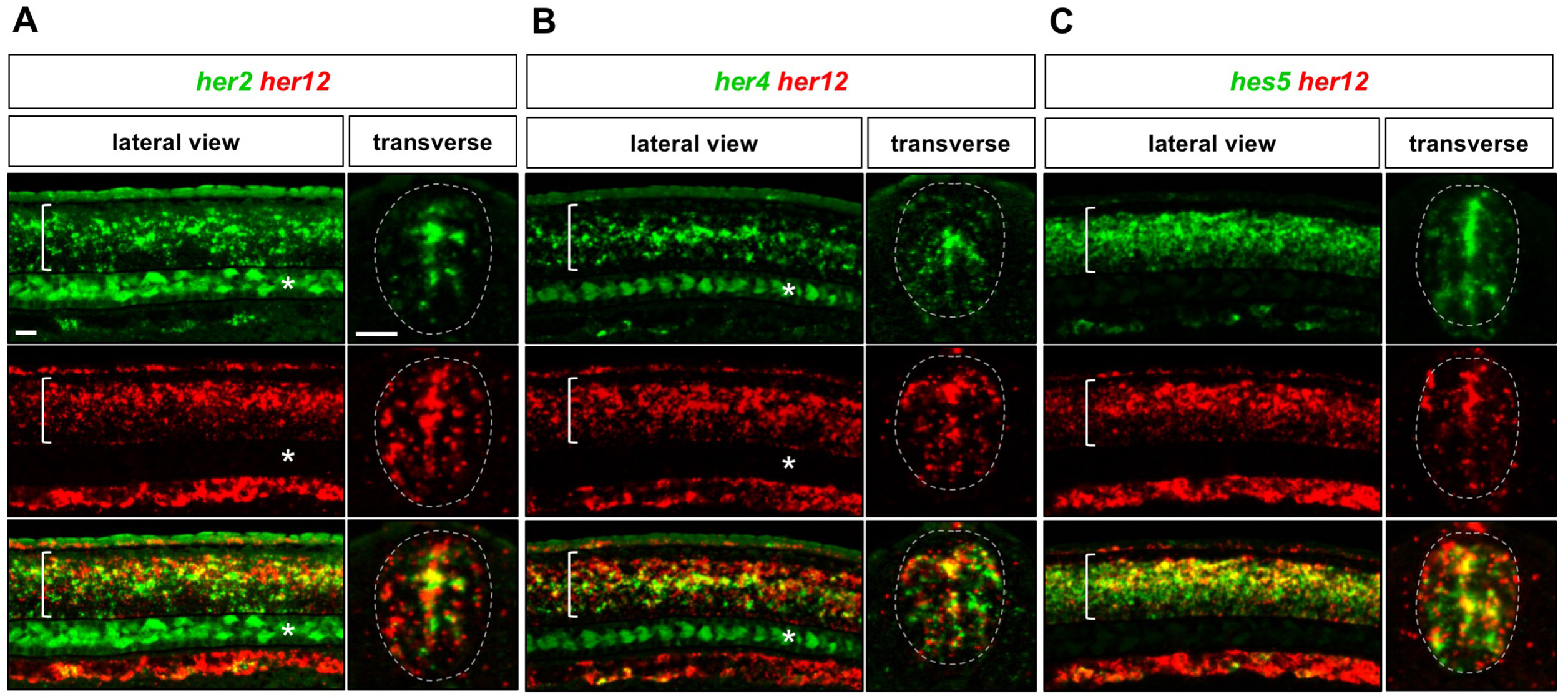
Co-expression of *her12* and other Notch target genes in the spinal cord. Whole-mount double fluorescent in situ hybridisation was performed in wild-type embryos at 24 hpf for (A) *her2* and *her12*, (B) *her4* and *her12* and (C) *hes5* and *her12.* Brackets in lateral views and dotted lines in transverse views denote the extent of the spinal cord. Note that *her2* and *her4* expression in notochord cells is indicated by asterisks. Scale bars: 20 μm.

**Figure S2.**
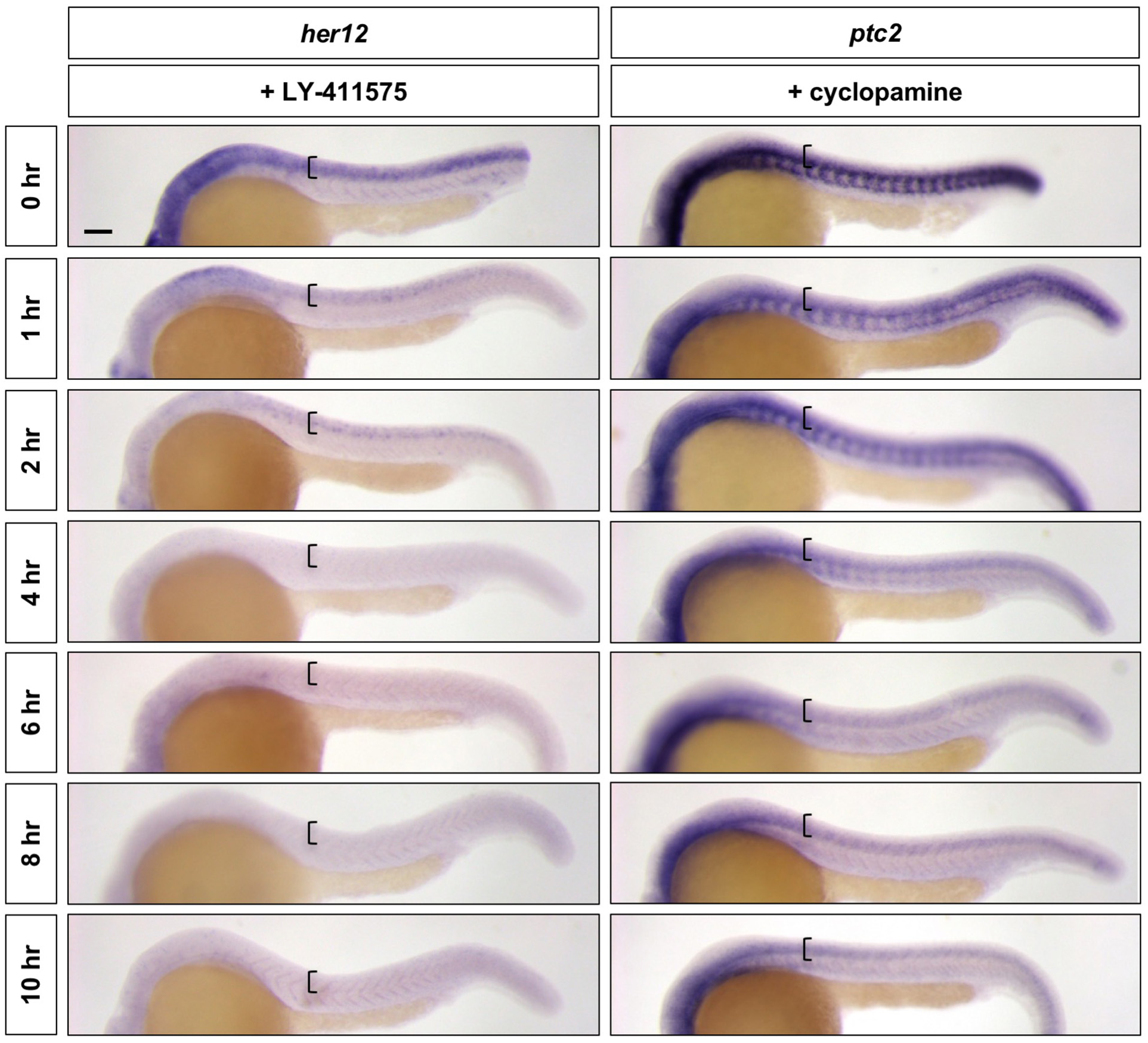
LY-411575 and cyclopamine are efficient inhibitors of Notch and Hh signalling, respectively. Wild-type embryos were treated with LY-411575 or cyclopamine for 0, 1, 2, 4, 6, 8, or 10 hours, and fixed at 24 hpf. Whole mount in situ hybridisation was then performed for *her12* in LY-411575 treated embryos and *ptc2* in cyclopamine treated embryos. Brackets denote the extent of the spinal cord. Scale bar: 20 μm.

**Figure S3.**
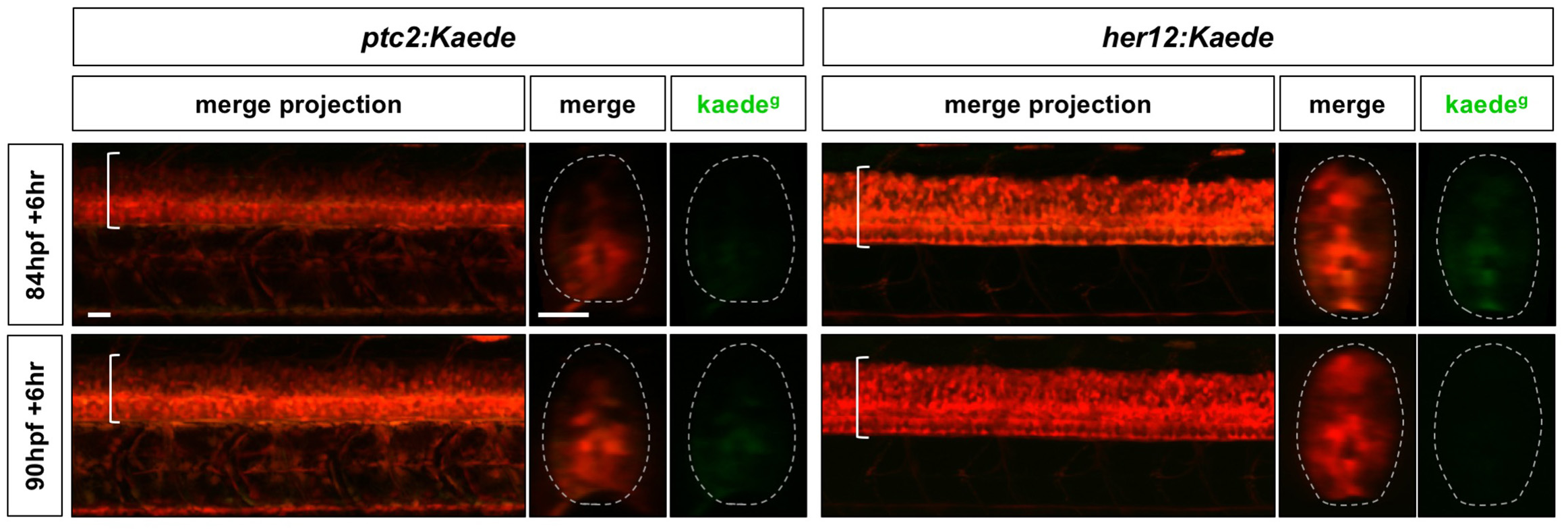
Notch and Hh response profile after the “signalling termination” phase. Continuation of the time course described in Figure 2, where *her12:Kaede* and *ptc2:Kaede* embryos were photoconverted at 84 hpf or 90 hpf, and imaged 6 hours post-conversion. Lateral views of confocal projections and transverse views of single slices are shown. Kaede^g^ panels show *de novo* synthesised Kaede^green^ after the time of photoconversion, while the merge panels show both previous Kaede^red^ expression and new Kaede^green^ expression. Brackets in lateral views and dotted lines in transverse views denote the extent of the spinal cord. Scale bars: 20 μm.

**Figure S4.**
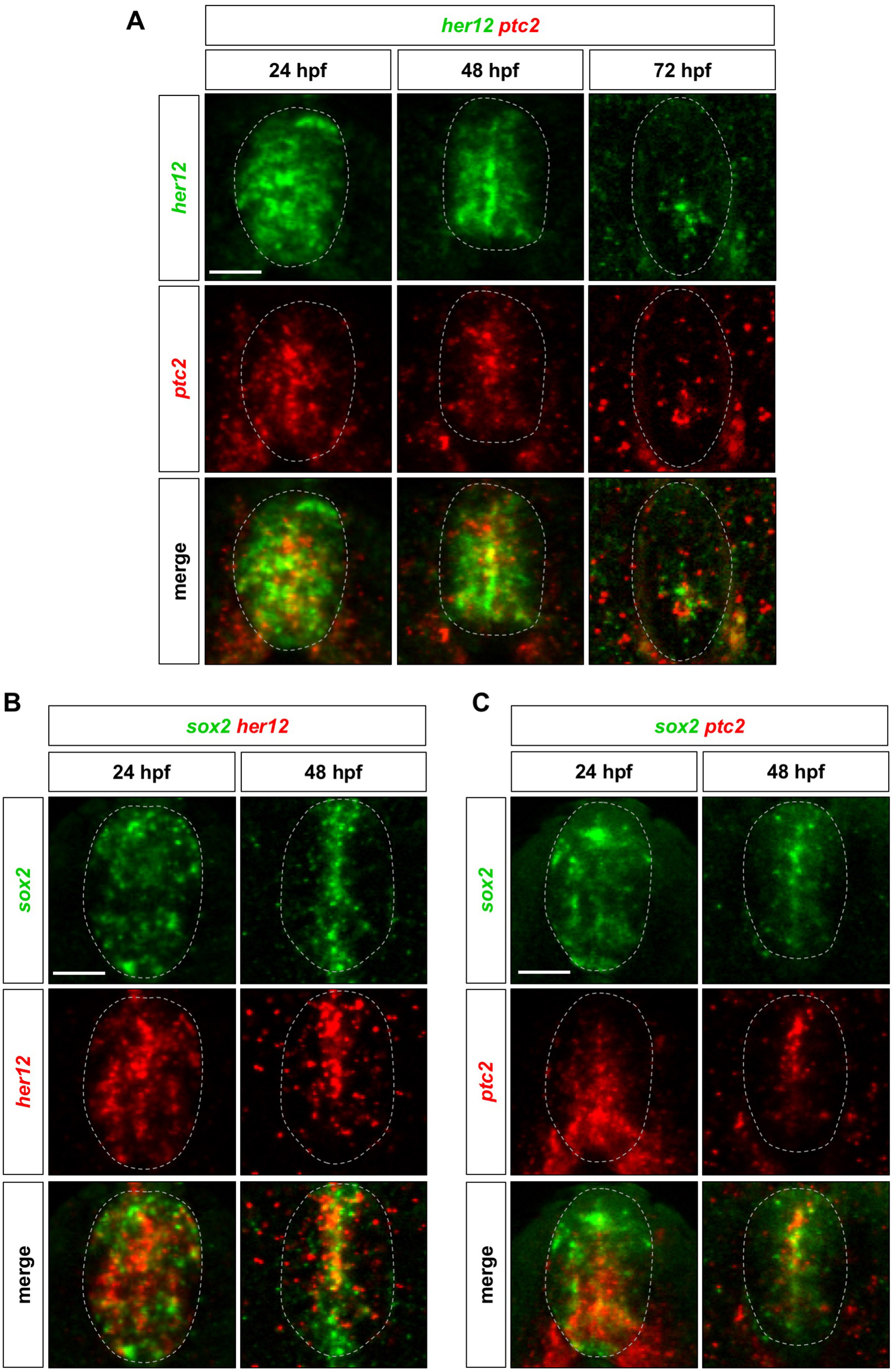
*her12* and *ptc2* are expressed in the same domain corresponding to *sox2^+^* neural progenitors. Whole-mount double fluorescent in situ hybridisation was performed in wild-type embryos for *her12* and *ptc2* (A), *sox2* and *her12* (B), and *sox2* and *ptc2* (C) at either 24, 48, or 72 hpf. Dotted lines denote the extent of the spinal cord. Scale bars: 20 μm.

**Figure S5.**
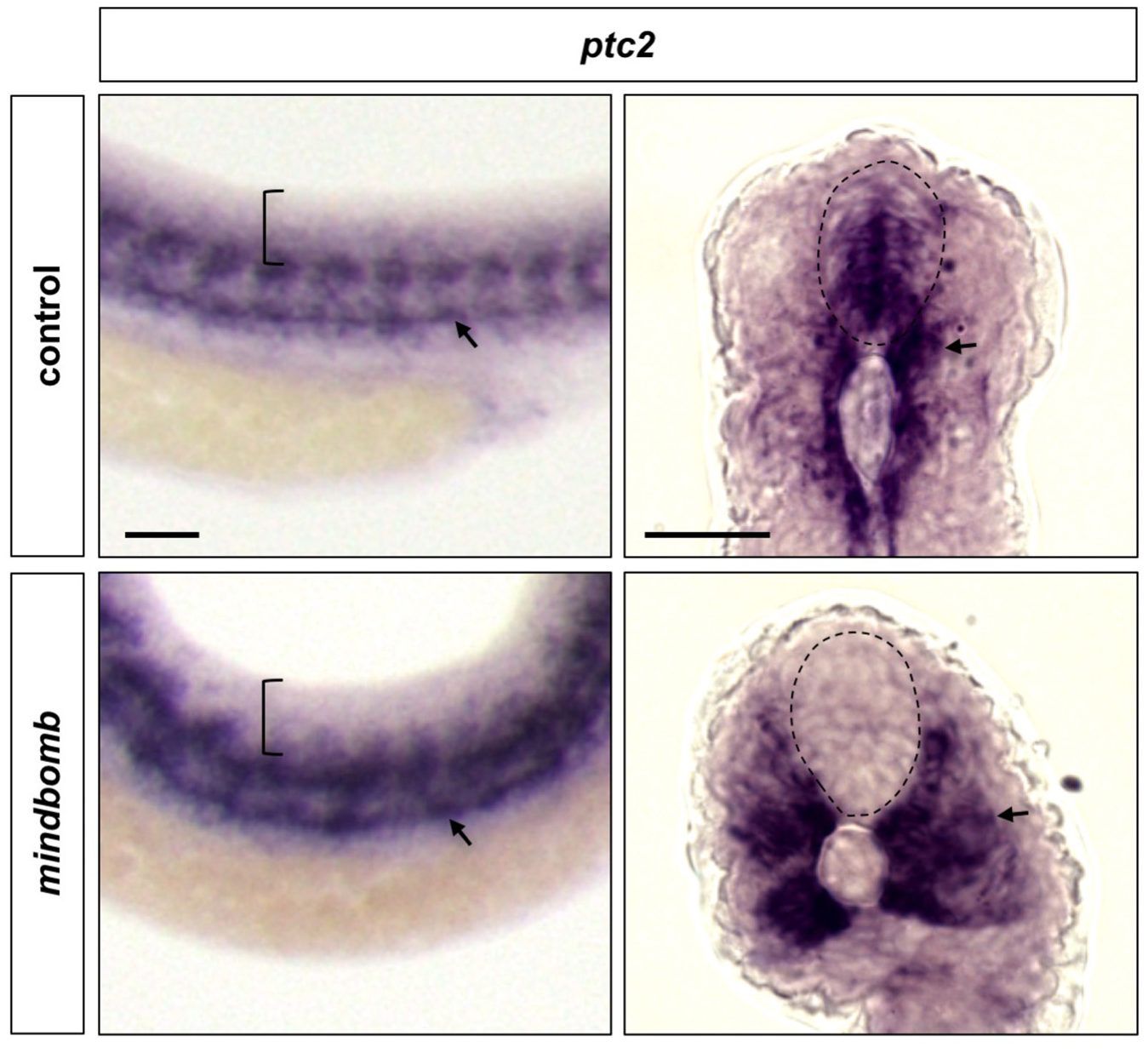
*mindbomb* mutants have no *ptc2* expression in the spinal cord. Whole-mount in situ hybridisation was performed in *mindbomb* mutants or their sibling controls for *ptc2* expression at 30 hpf. Dotted lines denote the extent of the spinal cord. Arrows indicate *ptc2* expression in somites. Scale bars: 20 μm.

**Figure S6.**
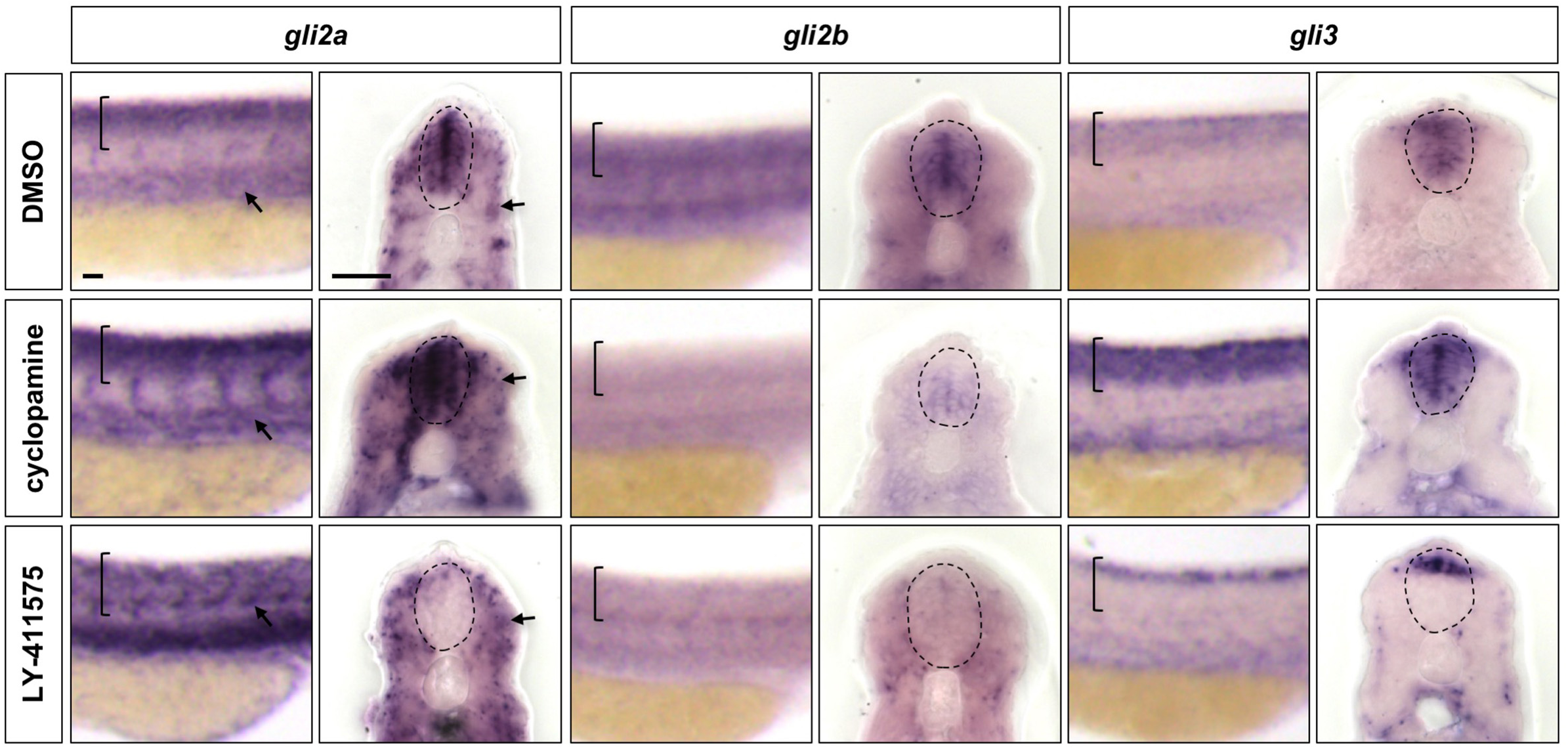
Notch signalling regulates the expression of all Gli family members in the spinal cord. Wild-type embryos were treated with DMSO, cyclopamine, or LY-411575 from 20 to 30 hpf, and stained with *gli2a*, *gli2b* or *gli3* probes. Brackets in lateral views and dotted lines in transverse views denote the extent of the spinal cord. Arrows indicate *gli2a* expression in somites. Note that in LY-411575 treated embryos, *gli3* expression is absent in most of the ventral spinal cord but is maintained in very dorsal region. Scale bars: 20 μm.

**Video S1. The re-emergence of *ptc2:Kaede* expression after photoconversion.** A *ptc2:Kaede* embryo was photoconverted at 28 hpf and then underwent time-lapse imaging for 18 hours. The vertical line indicates the boundary between photoconverted and unconverted regions at the start of the movie. Bracket indicates the extent of the spinal cord. Scale bar: 20 μm.

**Video S2. The *ptc2:Kaede* response profile along the anterior-posterior axis.** *ptc2:Kaede* embryos were photoconverted at 48 hpf and imaged 6 hours after. Individual transverse sections generated by 3D reconstruction were prepared into a video. The first frame is the most anterior slice and each subsequent frame moves further posterior through the embryo. The merge (left) and *Kaede^green^* (right) channels are shown. The spinal cord is denoted by solid lines and the active signalling domain (*Kaede^green^*) above the spinal canal is indicated by an arrowhead. Note that Figure 2C shows one single slice in the middle of the converted region. Scale bar: 20 μm.

**Video S3. The *her12:Kaede* response profile along the anterior-posterior axis.** *her12:Kaede* embryos were photoconverted at 48 hpf and imaged 6 hours after. Individual transverse sections generated by 3D reconstruction were prepared into a video. The first frame is the most anterior slice and each subsequent frame moves further posterior through the embryo. The merge (left) and *Kaede^green^* (right) channels are shown. The spinal cord is denoted by solid lines and the active signalling domain (*Kaede^green^*) above the spinal canal is indicated by an arrowhead. Note that Figure 2C shows one single slice in the middle of the converted region. Scale bar: 20 μm.

